# Gram-positive bacteria evade phage predation through endolysin-mediated L-form conversion

**DOI:** 10.1101/2022.05.24.493201

**Authors:** Jan C. Wohlfarth, Miki Feldmüller, Alissa Schneller, Samuel Kilcher, Marco Burkolter, Martin Pilhofer, Markus Schuppler, Martin J. Loessner

## Abstract

Bacteriophages kill bacteria by osmotic lysis towards the end of the lytic cycle. In the case of Gram-positive bacteria, peptidoglycan-degrading endolysins released at the end of infection cycle cause explosive cell lysis not only of the infected host, but can also attack non-infected bystander cells. Here, we show that in osmotically stabilized environments, *Listeria monocytogenes* can evade phage predation by transient conversion to a cell wall-deficient L-form state. This L-form escape is triggered by endolysins disintegrating the cell wall from without, leading to turgor-driven extrusion of wall-deficient, yet viable L-form cells. Remarkably, in absence of phage predation, we show that L-forms can quickly revert to the walled state. These findings suggest that L-form conversion represents a population-level persistence mechanism to evade complete eradication by phage attack. Importantly, we also demonstrate phage-mediated L-form switching of the urinary tract pathogen *Enterococcus faecalis* in human urine, which underscores that this escape route may be widespread and has important implications for phage- and endolysin-based therapeutic interventions.

## Introduction

In natural environments, bacteria are challenged by bacteriophages, which exert a permanent evolutionary pressure on microbial communities. In fact, phage-induced lysis is considered the most frequent cytocidal event in the biosphere^1^. The biology of phage infection has been subject to extensive studies and begins with phage attachment to the bacterial host surface by binding to a suitable receptor. In Gram-positive bacteria, host surface recognition typically involves carbohydrates that are covalently linked to the peptidoglycan cell wall, such as teichoic acids^2–4^. Receptor binding is essential and without it, infection cannot be initiated. After attachment, the phage genome is injected into the host followed by expression of viral genes and assembly of new virions. In order to release progeny, bacteriophages have to escape from their bacterial host cell. However, the cell wall and cytoplasmic membrane represent natural barriers preventing dissemination. Therefore, the tailed bacteriophages (*Caudovirales*) have evolved a canonical set of lysis proteins, designated the holin-endolysin system, which typically mediates host cell destruction by cell wall hydrolysis^5,6^. Endolysins are peptidoglycan hydrolases that specifically recognize and cleave the bacterial cell wall. The structure of these proteins is highly modular and typically consists of a N-terminal enzymatically active domain (EAD), and a C-terminal cell-wall binding domain (CBD) which promotes substrate specificity^7–9^. Access of endolysins to their substrate must be tightly regulated and depends on the assembly of holins in the cytoplasmic membrane at the end of the lytic cycle. This leads to pore formation, membrane depolarization, and access of endolysin to the cell wall, facilitating immediate degradation the peptidoglycan^5,7,10^. Together, these effects result in explosive cell lysis of the host^8,11,12^.

Remarkably, recent studies have demonstrated that phage-induced lysis may concomitantly also result in a massive release of bacterial membrane vesicles (MV) from both Gram-positive and Gram-negative bacterial cells^12,13^. These MVs incorporate cytosolic content including genomic DNA, thus sharing some similarity with cell wall-deficient L-form cells^11^. However, the potential role of L-form switching in the natural interaction of bacteria and their phage predators has not been established. The available evidence shows that many bacteria may transiently enter a wall-deficient state in the presence of certain triggers, such as lytic enzymes or cell wall-active antibiotics ^14,15^. Due to the lack of a cell wall and associated molecules, L-forms are intrinsically resistant to such peptidoglycan-targeting compounds. Therefore, L-form research has mostly addressed their possible role as persisters in chronic or recurrent infections^16–19^. While it has been pointed out long ago that L-forms may also confer resistance to phage infection^20,21^, the biological relevance of this phenomenon remained unclear, because no clear link between phage infection and L-form emergence could yet be demonstrated.

Here, we investigate the effects of phage infection on the emergence of bacterial L-forms, using *L. monocytogenes* and phage A006. We have recently developed a model for studying the biology of transient *Listeria monocytogenes* L-forms. These cells undergo an efficient L-form switch in the presence of an inducer, such as penicillin or lysozyme, while retaining the ability to revert to the walled state in absence of selective pressure^22,23^. Remarkably, L-form proliferation neither requires a cell wall nor the FtsZ-based cell division machinery^24^. In contrast, proliferation seems to rely solely on continuous membrane synthesis and biophysical effects, where an increased surface area-to-volume ratio results in membrane protrusion and formation of internal or external vesicles as viable progeny^25,26^.

Temperate *Listeria* phage A006 is a member of the *Siphoviridae* and comprises a 38.1 kB double-stranded DNA genome^27^. We have recently developed an L-form-based platform that allows facile and rapid genetic manipulation of this phage. Due to its genetic tractability, it has emerged as a model to study *L. monocytogenes* phage-host interactions^22,28,29^.

We show that the presence of phages can trigger L-form conversion in bacterial populations, which confers resistance to further infection, and demonstrate that L-form conversion is also possible based on the activity of endolysin released during repeated cycles of phage infection. Mechanistically, endolysins act by inducing lesions in the cell wall, thereby promoting turgor-driven extrusion of wall-deficient cells. Importantly, phage-induced L-forms retain the ability to revert to the walled state in the absence of selective pressure. These effects are not restricted to *L. monocytogenes* but could also be observed in *Enterococcus faecalis* phage-host pairs. Together, our results suggest that Gram-positive bacteria can evade phage predation on the population level by a transient switch of subpopulations to the L-form state.

## Results

### Bacteriophage infection promotes release of wall-deficient cells under osmoprotective conditions

We initiated the current study by exploring the effect of virulent phage infection on the emergence of bacterial L-forms. These experiments were inspired by earlier observations that prophage-triggered cell lysis results in the emergence of bacterial membrane vesicles^13^. In principle, these vesicles comprise the minimum characteristics of cellular life including genomic DNA, cytosolic content and a cellular membrane^11^, thereby resembling L-forms. However, previous work was performed in hypotonic environments, thus preventing the emergence of L-form cells due to osmotic cell lysis^12,13^. We therefore aimed to induce phage-induced L-form switching under osmoprotective conditions. We used *L. monocytogenes* strain EGD-e Rev2, which can undergo efficient L-form switching and reversion under variable selective conditions^22^. To ensure a strictly lytic bacteriophage phenotype, we used A006 ΔLCR, an engineered virulent derivative of temperate phage A006 that lack its entire lysogeny control region^28^. To test our hypothesis, we developed a protocol in which we challenged Rev2 cells expressing chromosomally integrated eGFP with A006 ΔLCR in DM3 L-form medium containing succinate as an osmoprotectant and CaCl_2_ to support phage infection^30–32^. Under such conditions, phage-induced lysis still results in degradation of the thick peptidoglycan layer, while leaving the cytoplasmic membrane structurally intact (Fig. 1 and Supplementary Video 1). This leads to the formation of wall-deficient cells that remain stable even after prolonged incubation periods (Fig. 1a, Supplementary Video 1). In contrast, phage infection in standard hypotonic medium (0.5 BHI) dramatically decreased the half-life of wall-deficient cells and resulted in rapid osmotic lysis (Fig. 1c). As expected, exposure to the parental temperate phage A006 yielded similar results, thus demonstrating that emergence of wall-deficient cells also occurs after infection with wild-type temperate phage during its lytic reproduction cycle (Fig. 1b).

**Fig. 1:**
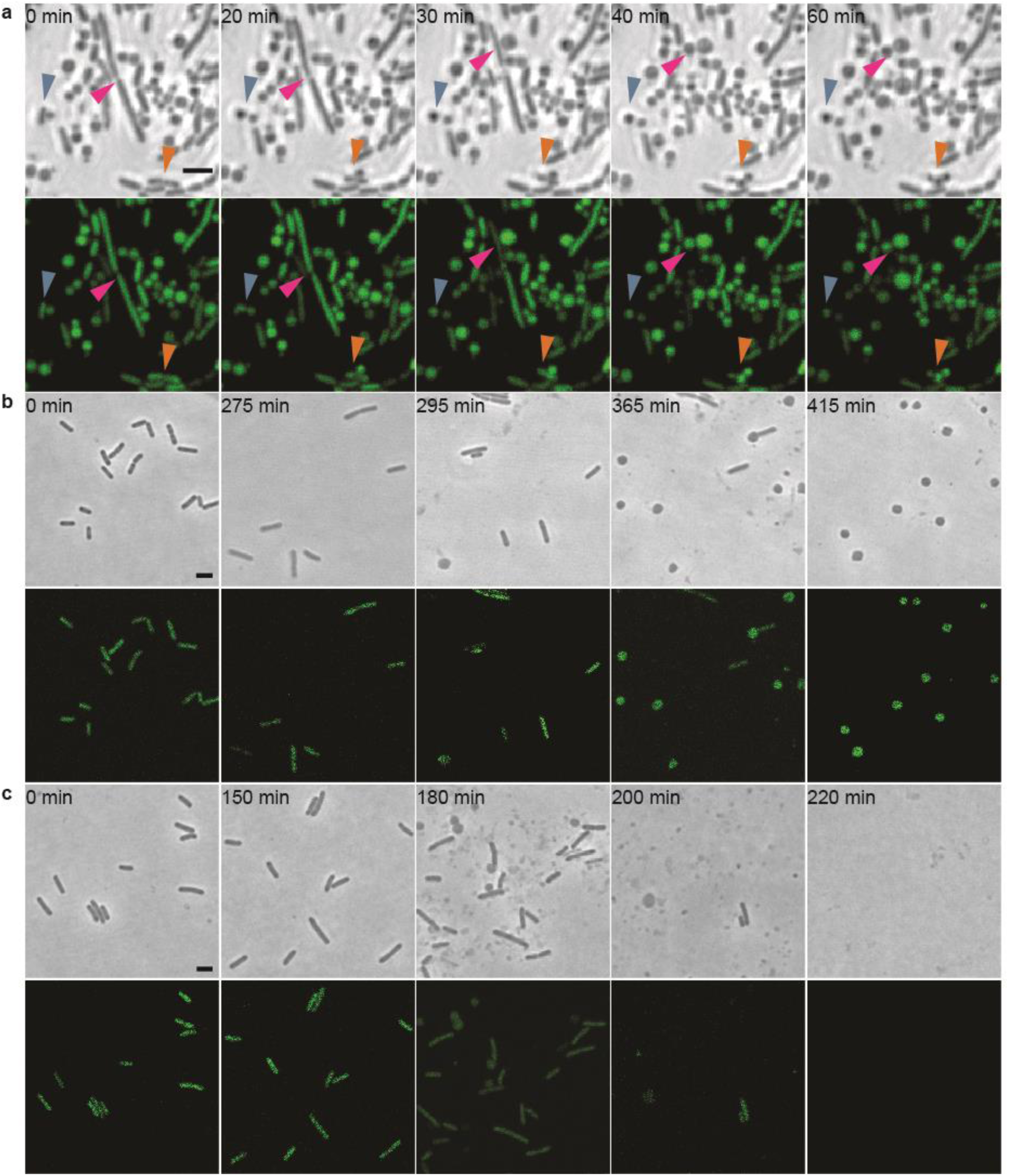
Phage-induced lysis leads to formation of wall-deficient bacterial cells under osmoprotective conditions. a, The effects of infection with a lytic phage on the transition of *L. monocytogenes* Rev2 to wall-deficient bacterial cells under osmoprotective conditions. Cells expressing chromosomally integrated eGFP were challenged with a strictly lytic mutant of temperate *Listeria* phage A006 lacking the lysogeny control region (ΔLCR) in osmoprotective DM3_Φ_ medium. Turbidity was monitored by spectrophotometry and before lysis, cells were placed on DM3 agar for time-lapse microscopy (t=0 min). Shown are phase contrast (PC) micrographs and the corresponding channel for green light emission. Arrows denote individual bacterial transition events. Individual frames are extracted from Supplementary Video 1. b-c, Contrary effects of hypotonic vs. osmoprotective medium on phage-induced wall-deficient cells. *Listeria monocytogenes* Rev2 cells expressing chromosomally integrated eGFP were infected with temperate phage A006 in osmoprotective DM3_Φ_ medium (b) or standard hypotonic 0.5 BHI medium (c). Cells were sampled at several timepoints before and after phage-induced bacterial lysis, followed by microscopic examination on 0.5 BHI or DM3 agar pads, respectively. In hypotonic medium, transition to the wall-deficient state is accompanied by rapid lysis, whereas under osmoprotective conditions, transition to the wall-deficient state is observed and most cells remain intact. Shown are PC images and the corresponding channel for green light emission, respectively. The figures are representative of at least three independent experiments. Scale bars, 2 µm.

### Phage infection triggers L-form switching and proliferation

Following the above observations, we asked whether the wall-deficient vesicles observed in Fig. 1 in fact represented viable L-forms. A hallmark for L-form cells and a distinction from protoplasts is their ability to proliferate in the absence of a cell wall^33^. Therefore, we aimed to observe proliferation using time-lapse microscopy. Towards this end, Rev2 cells expressing eGFP were infected with phage A006 ΔLCR and incubated for 18h to minimize the number of potential walled survivors that would overgrow the slow-growing L-forms. The culture was then transferred on osmoprotective agar for time-lapse microscopy, which demonstrated abundant wall-deficient cells undergoing shape deformations and irregular cell divisions characteristic for L-form growth^26^ (Supplementary Fig. 1 and Supplementary Video 2). To confirm and quantify the observed effect for wild-type phage A006 (Fig. 4g), we infected Rev2 cells with serial dilutions of phage at t=0min and monitored the infection dynamics by time-course turbidity assays and plating of lysed cultures on DM3 agar at various timepoints (Fig. 2a, b). Curiously, we found that the fraction of L-form survivors increased with decreasing phage concentration (Fig. 2b). At high phage concentrations, when most bacteria should be infected during the first cycle, bacterial survivors were predominantly walled. In contrast, infections at lower phage concentrations predominantly resulted in L-form colonies, which were phenotypically discernible by their characteristic “fried-egg” colony morphology (Fig. 2c, d). To explore whether these observations also hold true for other phages and bacterial species, we challenged *L. monocytogenes* Rev2 with several different phages including P35, P40, A118 (*Siphoviridae*), and A511 (*Myoviridae*). Moreover, to provide proof of principle for other Gram-positive bacteria, we challenged *Enterococcus faecalis*, which has also been reported to convert to L-forms^34^, with the virulent *Enterococcus* phage Efs7 (*Siphoviridae*) (Fig. 4h). Again, we observed emergence of L-form colonies for all tested combinations of phages and host strains (Fig. 2f, g). Importantly, both *L. monocytogenes* and *E. faecalis* L-forms retained the ability to switch back to the walled state, indicating that removal of selective pressure allows for reversion to the walled phenotype (Fig. 2e, h, i).

**Fig. 2:**
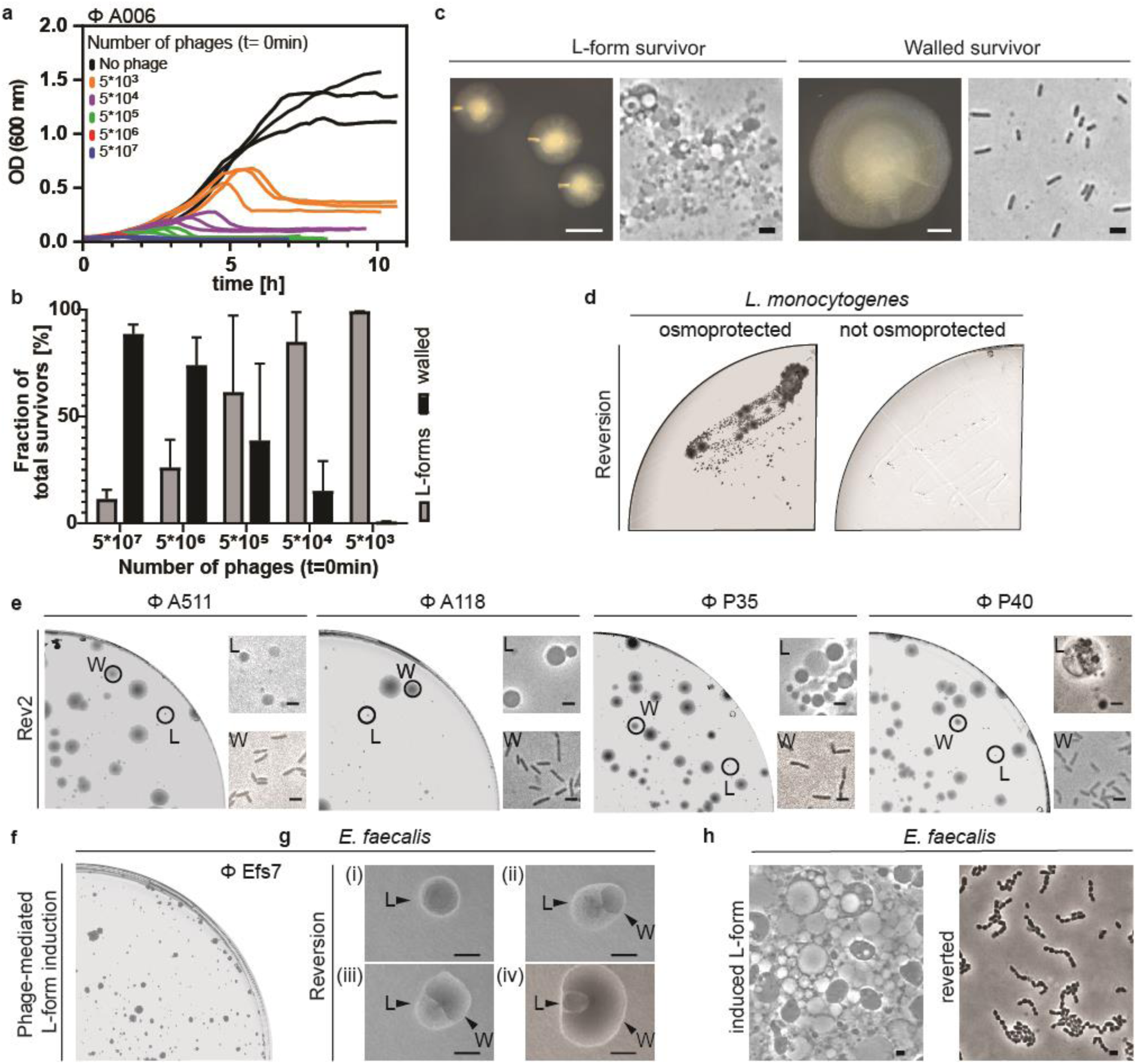
Phage infection triggers bacterial L-form switching. a-c, L-form switching of *L. monocytogenes* Rev2 in response to phage A006 infection at varying phage concentrations. a, Growth curves in liquid culture for *L. monocytogenes* Rev2. Bacteria were challenged with serial dilutions of phage (5*10^7^, 5*10^6^, 5*10^5^, 5*10^4^, 5*10^3^ PFU) or no phage at t=0min. For each dilution, three independent replicates are presented as individual curves. b, Relative quantification of bacterial colony forming units (CFU) on DM3 agar. L-form survivors and walled survivors are normalized to the number of the total CFU per infection. Values are displayed as mean of three biological replicates. Each culture was plated on osmoprotective DM3 agar at after approximately 5h, after reaching highest optical density (OD600 nm). Colonies of walled bacteria were enumerated after 2 days, L-form colonies after 5 days. c, Representative images of colony phenotypes and corresponding PC micrographs of *L. monocytogenes* Rev2 L-forms (left) and walled cells (right panel) on DM3 agar as observed in (b). d, Phage-induced *L. monocytogenes* Rev2 L-forms retain the ability to revert to the walled state in absence of selective pressure. Shown is a re-streak of a single survivor L-form colony obtained from (b) on osmoprotective DM3 agar and hypotonic 0.5 BHI agar. Note that numerous reverted walled colonies as well as non-reverted L-form colonies emerge on DM3 agar plates after 5d. In contrast, no colonies emerge on 0.5x BHI agar, ruling out the possible presence of remnant walled cells in the analyzed colonies. e, Infection of *L. monocytogenes* Rev2 with phage A511, A118, P35 or P40 triggers L-form switching. As above, liquid cultures were infected at different phage concentrations and streaked on DM3 agar after lysis. Plates were imaged 5d post-infection and are representative of individual infections at 510^3^ PFU (A511, A118, P40) or 5*10^6^ PFU (P35). Black circles mark representative L-form (L) or walled (W) colony phenotypes. Images of phase-contrast (PC) microscopy are shown for each colony. f, Infection of *E. faecalis* with phage Efs7 triggers L-form conversion. Liquid cultures of bacteria were infected with 10^3^ PFU and streaked on DM3 agar supplemented with 3.2 mM L-cysteine after lysis. Plates were imaged 2d post infection. g, *E. faecalis* L-forms retain the ability to revert to the walled state. Shown are different L-form colonies at different stages of reversion (i-iv) as observed 2d post infection. L-form colonies can be easily discerned from their walled counterparts due to different refraction properties. h, PC micrographs show *E. faecalis* L-forms and reverted bacteria. Scale bars, 0.5 mm (c, left; g), 2 µm (c, right; e; h).

### L-form escape is mediated by collateral damage from liberated endolysins

The observation of phage-induced L-form conversion raised the question regarding its primary effector. It has been shown that phage lysis may trigger formation of bacterial membrane vesicles^12,13^. Therefore, we speculated that the concomitant release and accumulation of endolysins following repeated infection cycles could be involved in L-form generation. This idea was supported by our observation that infections at low initial phage concentrations increase the fraction of L-form survivors compared to higher concentrations of applied phage (Fig. 2c). To investigate the effect of endolysins on L-form emergence, we produced recombinant phage A006-derived endolysin Ply006^35^ and Efs7-derived endolysin Ply007 (C-terminally linked to a 6xHis-Tag) (Fig. 3f). Next, we challenged intact *L. monocytogenes* Rev2 and *E. faecalis* cells with serial dilutions of the respective purified endolysin (Fig. 3a, b). Based on the linear ranges of the enzymes in DM3_Φ_ we determined a specific activity of approximately 0.4 ΔOD(600 nm min^- 1^ µM^-1^) for both Ply006 and Ply007 (Fig. 3c), demonstrating the high activity of the enzymes on their specific cell wall substrate. To test whether endolysin-mediated lysis promotes L-form generation, we then plated lysed bacterial cultures on osmoprotective DM3 agar. Indeed, L-form colonies could be observed at high frequency (approx.1-2 % of lysed cells) after 2-5 d incubation for both *E. faecalis* and Rev2 (Fig. 3d, e). These results showed that endolysins can act as efficient “transforming agents” for L-form conversion.

**Fig. 3.**
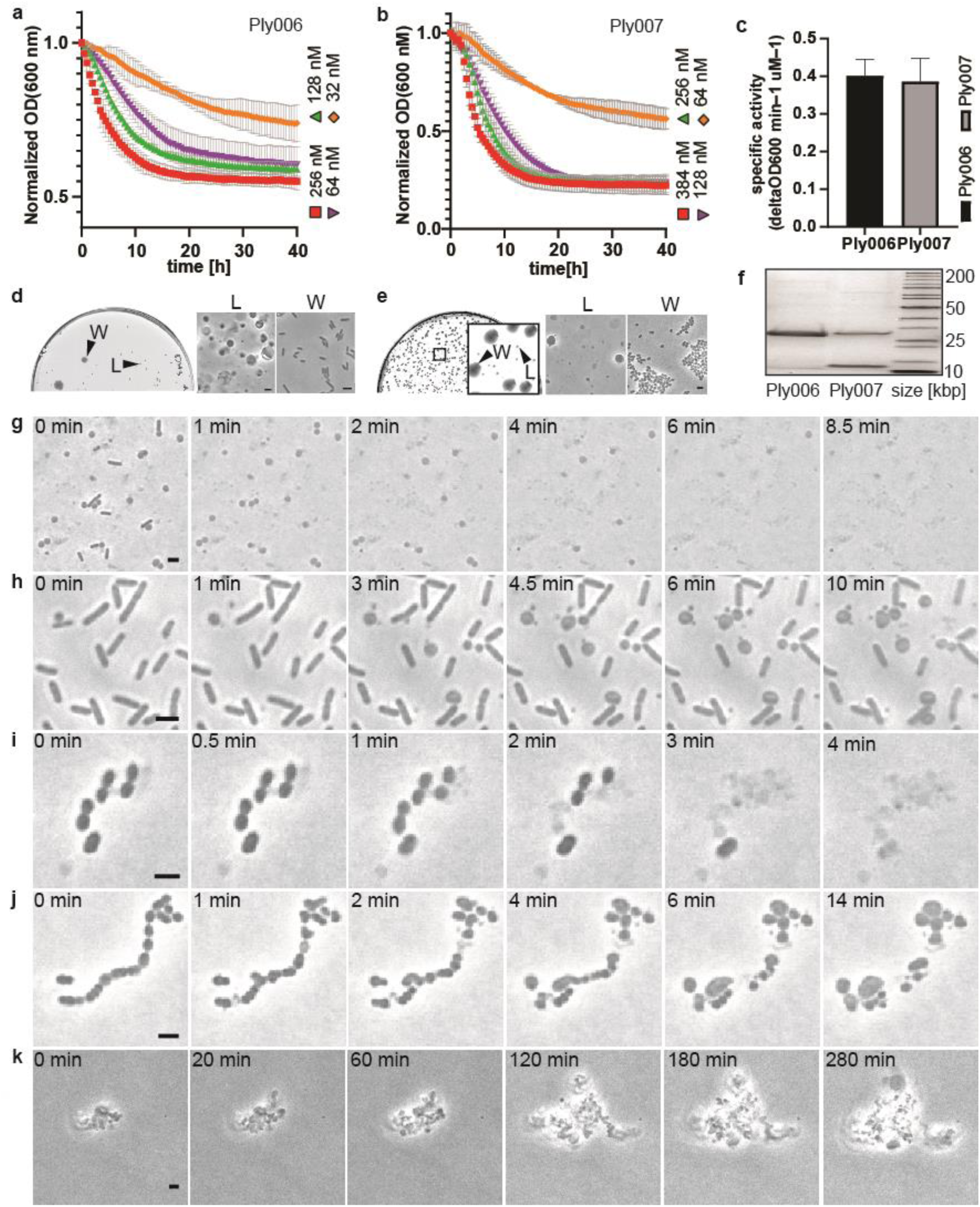
Endolysin activity promotes L-form conversion under osmoprotective conditions. a-b, Turbidity reduction of *L. monocytogenes* Rev2 and *E. faecalis* in liquid medium, upon exposure to serial dilutions of Ply006 or Ply007, respectively. Data are normalized to OD(600 nm) at t=0 min, and error bars reflect standard deviation of mean of three independent experiments (n=3). c, Specific enzyme activities of Ply006 and Ply007 expressed as ΔOD(600 nm min^−1^ μM^−1^) were calculated from the linear activity range of the enzymes in (a) and (b). d-e, Endolysins trigger bacterial L-form generation. d, The effects of Ply006 on *L. monocytogenes* Rev2 L-form emergence. Cells were exposed to 1 μM Ply006 for 40 min. Aliquots of 50 µl were streaked on DM3 agar in 10x serial dilutions and incubated for 5d. Shown is a plate at 10^−4^ dilution with clearly discernible walled and L-form colony morphologies. PC micrographs were taken from the plate shown on the left. e, The effects of Ply007 on *E. faecalis* L-form emergence. Cells were exposed to 1 μM Ply007 for 40 min. Aliquots of 50 µl were plated on DM3 agar supplemented with 3.2 mM cysteine in 10x serial dilutions and incubated for 2d. Shown is a plate at 10^−3^ dilution with clearly discernible walled and L-form colony morphologies. PC micrographs were taken from the plate shown left. f, SDS-PAGE of purified Ply006, 25.7 kDa and C-terminally 6xHis-tagged Ply007 big subunit, 28.2 kDa and corresponding 6xHis-tagged small subunit, 10.01 kDa, see also **Fehler! Verweisquelle konnte nicht gefunden werden**.. Samples of 2 μg protein were loaded per lane. g-j, The effects of endolysin on bacteria under hypotonic or osmoprotective conditions. g-h, The effects of Ply006 on *L. monocytogenes* Rev2 cells under hypotonic (g) or osmoprotective (h) conditions. Cells expressing chromosomally integrated eGFP were challenged with 1 μM Ply006 and immediately transferred on standard hypotonic 0.5 BHI agar or osmoprotective DM3 agar for time-lapse microscopy. Individual PC micrographs are extracted from Supplementary Videos 3 and 4a. i-j, The effects of Ply007 on *E. faecalis* cells under hypotonic (i) or osmoprotective (j) conditions. Cells were exposed to 1 μM Ply007 and immediately transferred on standard hypotonic 0.5 BHI agar or osmoprotective DM3 agar supplemented with 3.2 mM cysteine for time-lapse microscopy. Individual PC micrographs are extracted from Supplementary Videos 5 and 6. k, Time-lapse of Ply007 induced *E. faecalis* L-form proliferation. PC micrographs are extracted from Supplementary Video 7. Scale bars, 2 µm (b, d-k).

### Endolysins enable L-form generation by inducing localized lesions in the cell wall

To get a mechanistic insight into endolysin mediated L-form switching, we exposed walled *L. monocytogenes* Rev2 cells expressing eGFP or *E. faecalis* cells to 1 µM Ply006 or Ply007, respectively, and followed L-form escape via single-cell resolution time-lapse microscopy. We observed that under osmoprotective conditions, endolysin-mediated L-form conversion typically started by a blebbing process, resulting in extrusion of the cytoplasmic membrane from the cell wall sacculus, followed by proliferation of the wall-deficient cells. In addition, we occasionally observed transition events following explosive cell lysis (Fig. 3h, j, k; Supplementary Videos 4a, c; 6, 7). In contrast, endolysin treatment under hypotonic conditions usually leads to sudden osmotic rupture, disintegration of membrane vesicles, and cell death. This corroborates the initial finding that stability of phage-induced L-forms is dependent on osmoprotective environments (Fig. 3g, i; Supplementary Videos 3, 5). To investigate the ultrastructural underpinnings of endolysin-driven L-form conversion *in situ*, and in a near-native state, we employed cryo-electron tomography (cryoET). The diameter of intact *L. monocytogenes* or *E. faecalis* cells ranges from 600-800 nm, which is at the upper limits of sample thickness for conventional cryoET imaging^36^. To first test whether Rev2 and *E. faecalis* cells were suitable for imaging, we used cells that were directly plunge-frozen on EM grids. The obtained tomograms revealed clear visibility of all relevant bacterial structures including the cytoplasmic membrane and PG layer, confirming the technical feasibility of the approach (Fig. 4a, d). Next, we aimed to image L-form switching by inducing Rev2 and *E. faecalis* cells with 1 µM Ply006 or Ply007, respectively, followed by plunge-freezing. Indeed, tomograms of both *L. monocytogenes* Rev2 and *E. faecalis* showed the presence of many L-form-like cytoplasmic membrane vesicles (Fig. 4c, f). Further, we observed intermediate stages of membrane protrusions extruding through punctual lesions in the peptidoglycan cell wall (Fig. 4b, e). Based on multiple tomograms of cytoplasmic extrusions that were captured at different stages, we inferred that L-form switching comprises three distinct steps: (i) First, localized enzymatic hydrolysis causes the formation of punctures in the cell wall. Small membrane protrusions begin to extrude through these holes. (ii) Subsequently, the protrusions are filled with cytosolic content, driven by the internal turgor pressure of the cell. At this stage, the growing membrane bleb remains connected to the parental cell. iii) Finally, scission of the membrane bleb results in the formation of an independent and wall-deficient cell. Interestingly, we observed that Ply006-induced lesions in *L. monocytogenes* are preferentially located at the poles (Fig. 4b). This is consistent with previous studies demonstrating that cell wall binding domain of *Listeria* phage endolysin Ply006 and related enzymes preferentially attach to the polar regions of the cell wall^35,37^. Hence, it seems that the enzymatic function of Ply006 is spatially guided by its CBD. In contrast, no such site specificity was observed for the *Enterococcus* phage endolysin Ply007.

**Fig. 4:**
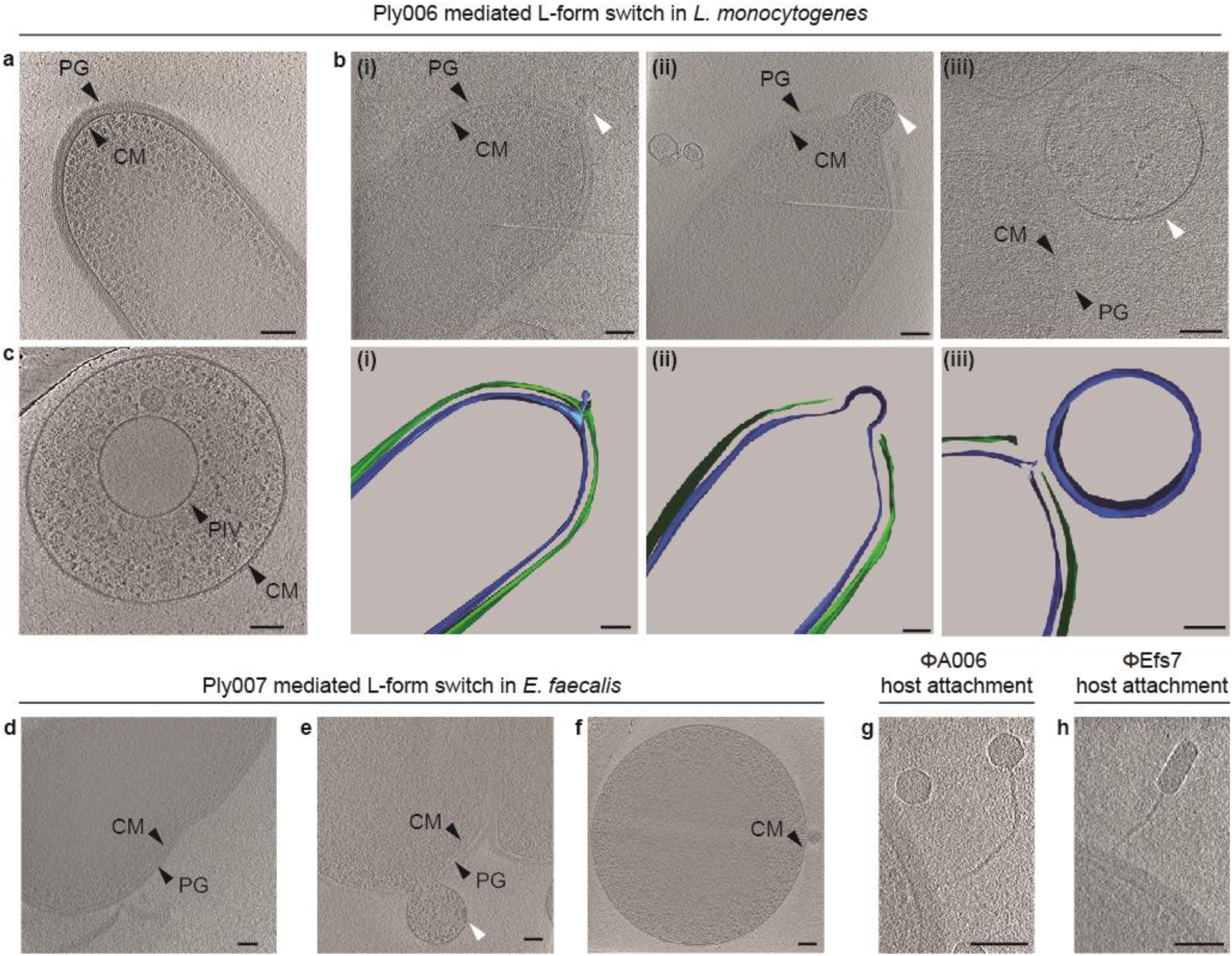
Endolysin-based disintegration of the cell wall induces L-form conversion. a-c, Cryotomography of *L. monocytogenes* Rev2 cells revealing the effects of endolysin Ply006 exposure on the bacterial cell wall *in situ*. a, Representative Rev2 walled cell with intact cytoplasmic membrane (CM) and peptidoglycan (PG) layer. Shown is a 26 nm-thick slice through a cryotomogram. b-c, Different stages of CM membrane blebbing (white arrows) in response to Ply006 exposure for 1 min (top), and corresponding segmented 3D model (bottom). Shown are representative images of CM-extrusions emerging from different cells. CM protrudes through lesions in the peptidoglycan layer, predominantly at the cellular poles. Blebbing occurs in different stages, ranging from small membrane protrusions (i) to blebs filled with cytoplasmic content (ii) and membrane-bound L-form like vesicles (iii). Shown are 15 nm-thick tomographic slices; PG: green, CM: blue. c, L-form like vesicle completely lacking detectable PG structures. A primary internal vesicle (PIV), located within the cell, is clearly visible. A 22 nm-thick tomographic slice is shown. d-f, Cryotomograms of *E. faecalis* Rev cells revealing the cell wall architecture *in situ* before (d) and after treatment with endolysin Ply007(e-f). d, Dividing *E. faecalis* coccus with a distinct PG layer and CM membrane. Shown is a 28 nm-thick tomographic slice. e-f, CM blebbing and induction of L-form like vesicles in response to Ply007 exposure. Shown are 22 nm-thick tomographic slices. g, Cryotomogram of a A006 virion attaching to a Rev2 host cell, 5 min post-infection (bottom). Shown is a 18 nm-thick tomographic slice. h, Cryotomogram of a Efs7 phage attachment to a *E. faecalis* Rev host cell, 10 min post infection. A 22 nm-thick tomographic slice is shown. PG, peptidoglycan layer; CM, cytoplasmic membrane. Scale bars, 100 nm. Also see Supplementary Videos 8 and 9.

### Cells undergoing phage infection are no longer capable of L-form escape

The above results suggested that phage-induced L-form conversion is triggered by endolysins. However, we had no means to directly assess whether this phenomenon occurred primarily with non-infected bystanders (lysis-from-without), or possibly also as a result of phage infection (lysis-from-within). Employing a recently developed synthetic phage engineering platform^22^, we created a A006-based reporter phage expressing a fluorescent protein, allowing direct monitoring of infected cells. In order to obtain high expression levels, a modified *gfp* gene was inserted and placed under control of the strong A006 major capsid protein promoter P_*cps*_^28,29^. The strategy for design of corresponding genome fragments for phage assembly is shown in Supplementary Fig. 5a and b, followed by rebooting in L-form cells^22^. Correct genome sequence was confirmed by DNA sequencing. Plaque phenotypes and phage concentration-dependent host killing were validated using soft agar overlay assays (Fig. 5c, d), and lysis kinetics monitored in liquid culture (Fig. 5e). As expected, engineered A006::*egfp*_*cps*_ showed similar lysis characteristics as the wild type, and phage-induced eGFP production became detectable at 45 min after infection (Fig. 5g). If L-form conversion of phage-infected cells would be possible, infection with A006:*egfp*_*cps*_ would results in eGFP labelled *L. monocytogenes* cells, visible even after L-form conversion. To test this, we infected Rev2 walled cells expressing chromosomally integrated RFP with excess amounts of A006::*egfp*_*cps*_, resulting in transient eGFP fluorescence in the majority of walled cells (Fig. 5f, g). Eventually, phage-induced lysis caused a sharp decrease in fluorescence, due to explosive cell death by sudden osmotic lysis of virtually all infected cells. In contrast, the red-fluorescent RFP expressing cells showed massive L-form switching instead of complete lysis. In conclusion, these results strongly suggest that not the phage-infected bacteria, but the non-infected bystander cells are responsible for L-form generation.

**Fig. 5:**
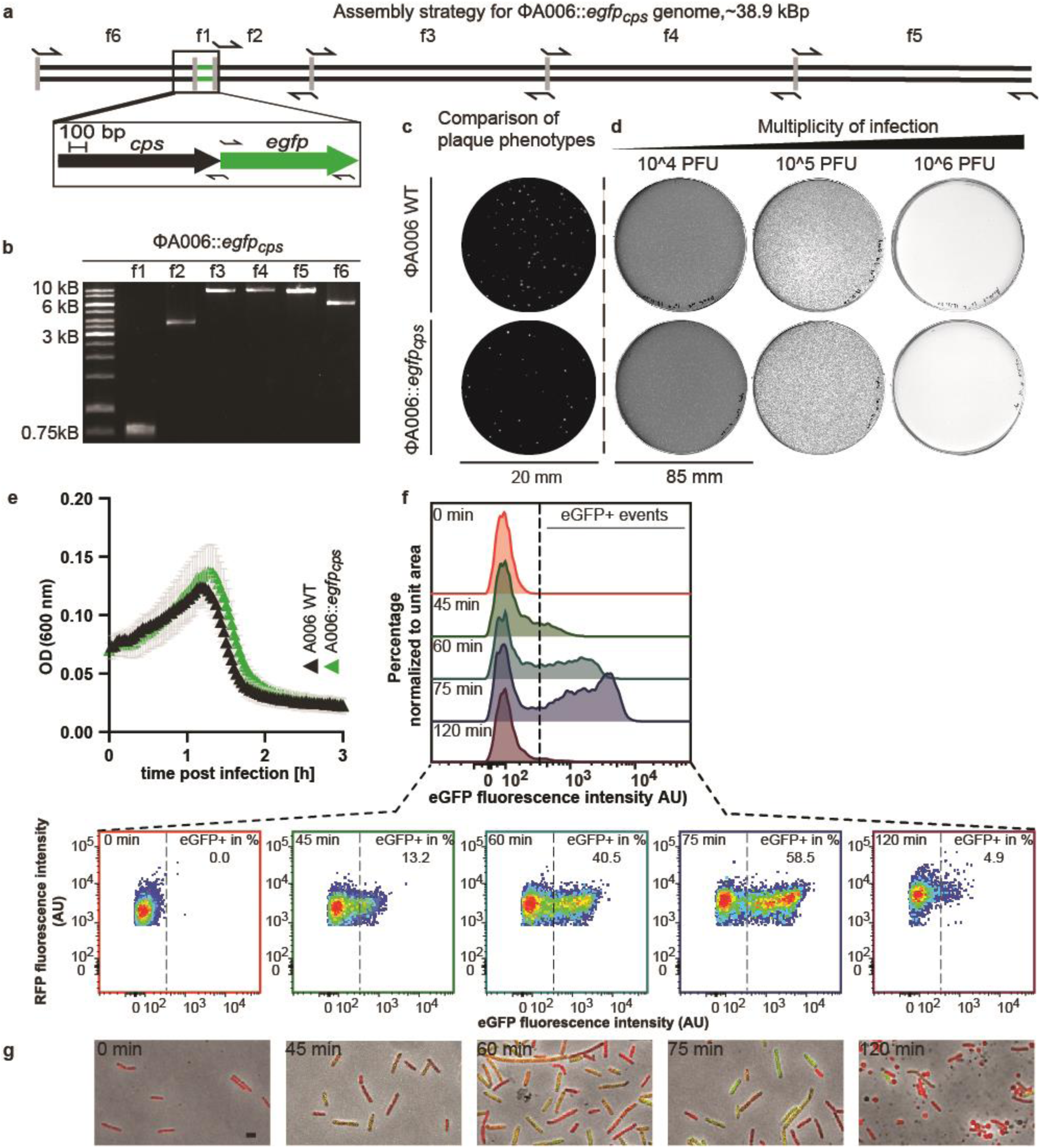
Bacterial cells undergoing productive phage infection do not form stable vesicles or wall-deficient L-form cells. a-b, Engineering of a phage that allows fluorescence-based single cell-tracking of productive phage infection in *L. monocytogenes*. Strategy for synthetic assembly of phage A006::*egfp*_*cps*_ (a) where *gfp* is placed under the control of the major capsid protein gene *cps* promotor, whose expression indicates productive phage infection. Arrows denote primer binding sites for PCR amplicons f1-f6 (b), which represent the basic segments for phage genome assembly. c, Comparison of plaque phenotypes between wild-type phage A006 and A006::*egfp*_*cps*_ as observed on soft agar overlay plates. Shown are representative plaque phenotypes 24 h post infection using *L. monocytogenes* Rev2 as a host (c-d). d, Concentration-dependent host killing in plate culture where phage A006 and its mutants are titrated in 10-fold serial dilutions (denoted as multiplicity of infection). e, Comparison of host killing kinetics in liquid infection (left panel) and corresponding fluorescence emission intensities (right panel). *L. monocytogenes* Rev2 bacteria were challenged with excess amounts of A006 or A006::*egfp*_*cps*_ in DM3_Φ_ liquid medium. Turbidity or fluorescence intensity was monitored for 3 h. Data are displayed as mean ± SD (n = 3). f, Flow cytometry analysis of *L. monocytogenes* Rev2 cells expressing chromosomally integrated RFP upon infection with excess amounts of A006::*egfp*_cps_ in DM3_Φ_ medium. Fluorescence-positive events (eGFP+) represent infected cells undergoing productive phage infection. Samples were analyzed at 0, 45, 60, 75, or 120 min post infection. Overlayed histograms depict eGFP fluorescence intensity (AU) vs. percentage of cells normalized to the mode. Pseudocolor density plots show eGFP fluorescence intensity (AU) vs. RFP fluorescence intensity (AU) displayed on a biexponential scale. The threshold value for eGFP+ events was set to 400 (AU, dashed black line). Data are representative of three independent biological experiments. g, Micrographs of bacterial cells taken from (f). Shown are merged images of channels for green and red-light emission, and PC channel. Walled bacterial cells undergo a strong “green-shift” due to transient phage-encoded GFP fluorescence. L-form switching occurs between t= 75 min and t= 120 min post infection. Note that wall-deficient cells (t=120 min) express RFP but not eGFP. Scale bar, 2 µm. AU, arbitrary units.

### Loss of cell wall-associated teichoic acids mediates phage resistance in *Listeria* L-forms

Phage infection of Gram-positive bacteria requires cell-wall associated binding ligands, such as wall teichoic acids covalently linked to the peptidoglycan of the host^2–4^. For example, the A006 receptor binding protein A006_gp17 is known to recognize specific sugar decorations of wall teichoic acids with high selectivity and sensitivity^38^. Because L-form conversion leads to a complete loss of the cell wall-associated phage receptors, it seemed reasonable to assume that L-forms are resistant to phage infection via this route. This idea was also supported by the massive emergence of L-forms observed here (Fig. 2). To provide formal proof, we exposed *L. monocytogenes* Rev2 L-forms expressing chromosomally integrated RFP to excess amounts of A006::*egfp*_*cps*_. Indeed, microscopic analysis revealed complete absence of fluorescence in L-forms even after prolonged periods of incubation, indicating that L-forms are not supporting phage binding and subsequent genome injection (Supplementary Fig. 3) due to a lack of wall teichoic acid ligands.

### Phage infection of *E. faecalis* in human urine leads to L-form emergence

Based on our observation of phage induced L-form switching, we asked whether this process may also be relevant under conditions found in a natural environment. Phage therapy is currently developed as a treatment option for several pathogens causing urinary tract infections, including Enterococcus faecalis^39–41^. It has recently been shown that urine provides the necessary osmoprotection to enable L-form switching and survival^17^. We therefore asked whether phage Efs7 infection of *E. faecalis* in human urine would potentially also result in L-form conversion. To test this hypothesis, we challenged the bacteria with serial dilutions of Efs7 in sterile-filtered human urine, followed by incubation overnight (Fig. 6a), plating, and quantification of the fraction of walled survivors and L-forms after two days (Fig. 6b-e). Strikingly, we found that Efs7 indeed induced a massive induction of L-forms in urine. In line with the results obtained for L. monocytogenes, excess amounts of phage reduce the fraction of L-form survivors after infection, whereas lower phage concentrations were more effective and resulted in L-forms as vast majority of bacterial survivors (Fig. 6b, c). Notably, almost all E. faecalis L-form colonies were able to undergo reversion to the walled state within 72 h (Fig. 6c).

**Fig. 6:**
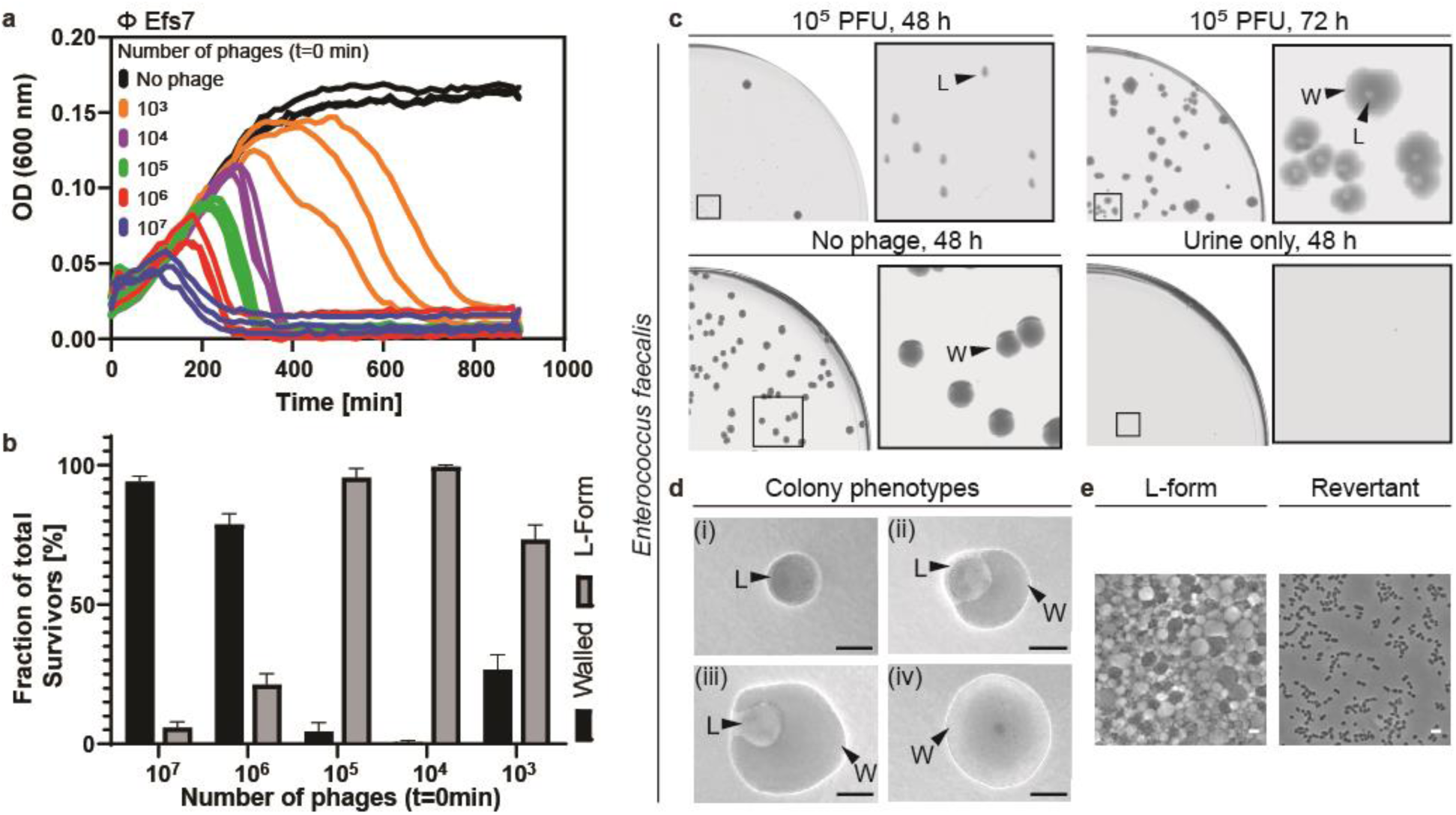
Phage-induced L-form conversion in human urine. a-c, Phage Efs7 infection of E. faecalis in human urine triggers L-form generation. a, Growth curves in human urine for E. faecalis. Bacteria were challenged with serial dilutions of phage (10^7^, 10^6^, 10^5^, 10^4^, 10^3^ PFU) or no phage at t=0min. As a control, sterile-filtered urine was incubated without bacteria (shown in (c) only). For each condition, three independent replicates are presented as individual curves. b-c, Relative quantification of L-form survival after phage infection. Cultures from (a) were plated 15h after infection on DM3 supplemented with cysteine. Populations of L-forms and walled survivors are normalized to the total CFU per infection. Quantification was performed 48h after streaking. Data are mean of three independent experiments. c-e, Effect of phage infection in human urine on L-form emergence and reversion. c, Representative high resolution plate scans obtained from (b). Top panel shows identical streaks of E. faecalis 48h and 72h after Efs7 infection with 10^5^ PFU. Note that after 48h all colonies show the L-form phenotype (L), followed by reversion back to the walled state (W) after 72h. Bottom panels show controls of E. faecalis with no phage or sterile-filtered urine only after 48h incubation. d-e, Individual colony phenotypes observed in (c). d, Micrographs of L-form colonies (i), intermediate states of reversion (ii, iii), and walled cells (iv). e, Phase-contrast micrographs are representative of walled E. faecalis cells as observed in revertant colonies and characteristic wall-deficient cells as observed in L-form colonies. Scale bars, 0.5 mm (d), 2 µm (e).

## Discussion

In the lytic cycle, phage infection normally results in sudden (sometimes explosive) lysis of the host, at least under standard culture conditions that are generally hypotonic. Here, we report that in an osmoprotective environment, Gram-positive bacteria such as *L. monocytogenes* or *E. faecalis* can manage to evade complete phage-induced lysis by switching to a transient wall-deficient L-form state, conferring resistance to further phage infection. Our results demonstrate that L-form conversion can occur due to collateral damage, which is cell wall hydrolysis from without caused by liberated, soluble phage endolysins. In consequence, this effect can contribute to transient persistence and rescue of viability of bacterial communities by enabling L-form escape of uninfected cells before phage infection can be initiated.

Based on our experiments with phage endolysins Ply006 and Ply007, we propose a mechanistic model of L-form escape that comprises three major steps: i) endolysin-mediated induction of punctual lesions in the cell wall and extrusion of small membrane protrusions ii) maturation, that is turgor-driven filling of the wall-deficient cell with cytosolic content including genomic DNA and iii) finally scission of cell membranes to form independent and viable L-form cells (Fig. 6).

**Fig. 6:**
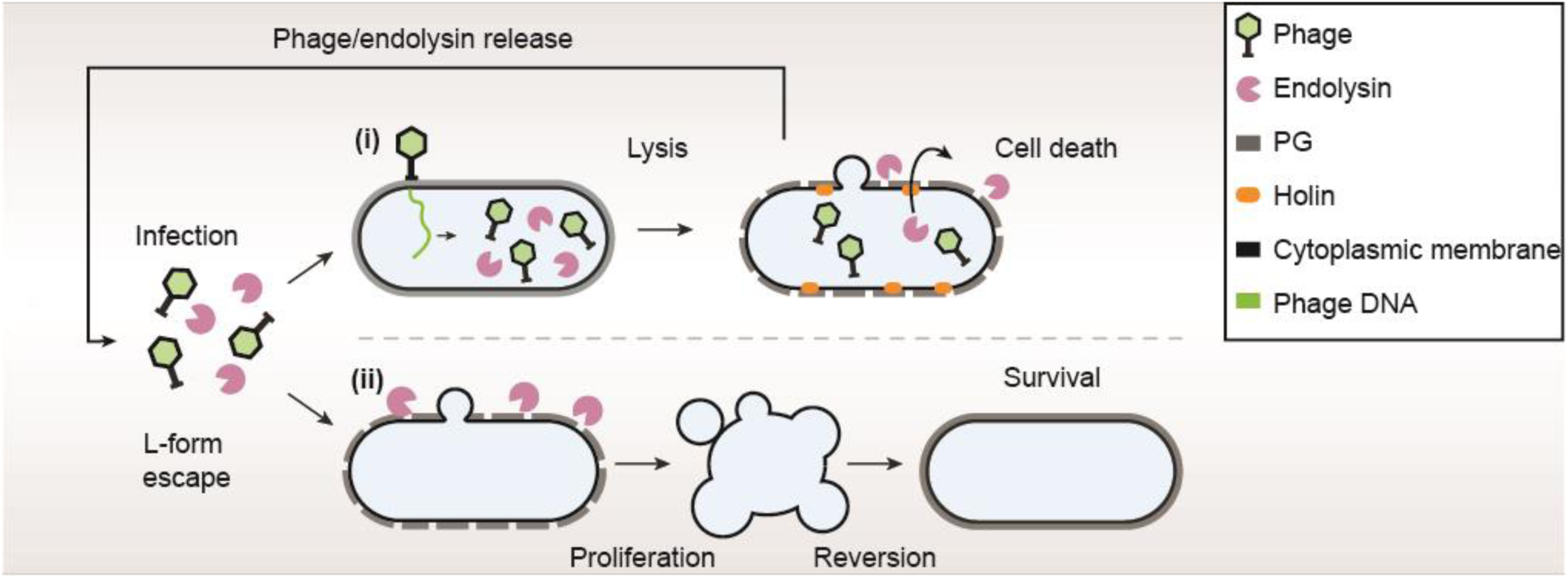
Model for phage-induced L-form escape as a survival strategy. The figure outlines two possible outcomes of an infection with a lytic bacteriophage in a bacterial population. At the time of lysis, infected cells release new phage as well as soluble endolysins. (i) Progeny phages may infect other bacterial cells resulting in lysis and cell death due to the concerted action of endolysins and holins. (ii) Endolysins promote L-form conversion by weakening the cell wall of so far uninfected, walled bystander cells. As a result, wall-deficient L-forms may occur that are completely resistant to phage infection while retaining the ability to proliferate. In absence of selective pressure, L-form cells are capable of reverting to the walled state.

Given that expression of endolysins at the end of the lytic cycle is a shared feature of all tailed phages, it is very likely that phage-induced L-form escape occurs among a wider range of Gram-positive bacteria, especially during growth in confined environments. This hypothesis is supported by our observation that L-form escape can be induced by different phages, including temperate and virulent members of the *Siphoviridae* and *Myoviridae*. Importantly, loss of the cell wall confers resistance of L-forms against viral infection, due to the lack of cell wall-associated phage receptors such as wall teichoic acids, and seems pivotal for L-form survival. Crucially, such phage-induced L-forms are only transiently wall-deficient and can revert to the walled state in absence of selective pressure. Therefore, this route may serve as a self-sustaining evasion mechanism to escape phage killing. Indeed, the massive emergence of L-forms following phage exposure indicates that endolysin-mediated L-form release is frequent and widespread, rather than an exceptional and singular event. Notably, due to very rapid enzyme kinetics, endolysin-mediated escape of phage-resistant L-forms occurs within seconds. Because bacteria often exist in dense communities, it is conceivable that endolysins released during lysis of phage-infected bacteria act on neighbouring cells even before they may be infected by progeny phage. This is consistent with our observation that the fraction of L-form survivors increases at decreasing number of applied phages.

Remarkably, this effect was also observed for uropathogenic *E. faecalis* in human urine as a suitable *ex vivo* environment, providing strong evidence that phage-induced L-form switching occurs during phage exposure of bacterial communities under natural conditions. Even though the impact and possible roles of L-forms in the environment remains elusive, nature provides a multitude of ecological niches that should in principle allow L-form growth. Noteworthy, L-forms have been previously reported to occur in a range of natural sources, including samples obtained from plants, animals and humans^16,18^. For example, a recent study demonstrated that L-forms may be frequently present in clinical urine samples of elderly patients^17^. In light of the phage-bacteria interactions uncovered in this study, in particular endolysin-mediated L-form escape and subsequent reversion to walled cells, this may have important implications for future efforts in the emerging field of phage and endolysin therapy. In fact, the generation of L-forms among Gram-positive pathogens could provide a source of bacterial survivors and persisters as a result of treatment with endolysins or replicating phages and may therefore be of particular importance and concern. This strongly emphasizes the need for application of additional effectors beyond the cell wall lytic activity of peptidoglycan hydrolases or phages, such as a combination treatment with non-cell wall active drugs and antibiotics.

## Materials and Methods

### Bacterial strains and growth conditions

Bacterial strains used in this study are listed in Table S1. Half-strength (0.5x) brain-heart infusion medium (BHI, Biolife Italiana) was used as a standard hypotonic medium for growth of *L. monocytogenes*, and BHI-FC (37g/L BHI, 4 g/L glycine, 6.06 g/L Tris, pH 7.5) was used as standard hypotonic medium for growth of *E. faecalis* at 30°C. L-forms were induced and grown in osmoprotective modified DM3 liquid medium, referred to as DM3_Φ_ (5 g/L tryptone, 5 g/L yeast extract, 0.01% BSA, 125 mM succinic acid, 180 mM glucose, 20 mM K_2_HPO_4_, 11 mM KH_2_PO_4_, 20 mM MgCl_2_, 200 uM CaCl_2,_ pH 7.3) at 30°C. DM3 agar (5 g/L tryptone, 5 g/L yeast extract, 0.01% BSA, 500 mM succinic acid, 180 mM glucose, 20 mM K_2_HPO_4_, 11 mM KH_2_PO_4_, 20 mM MgCl_2_, pH 7.3)^42^ was used for L-form growth on plate. For *E. faecalis* L-forms, DM3_Φ_ liquid medium and DM3 agar were supplemented with 3.2 mM L-cysteine. Microaerophilic conditions for growth in plate culture were generated using microaerophilic atmosphere generation bags (BioMerieux) in an anaerobic jar. *Escherichia coli (E. coli)* strains XL1 Blue MRF’ and BL21 Gold (DE3) were grown in LB medium (10 g/L Tryptone, 5 g/L Yeast extract, 5 g/L NaCl) at 37°C. If required, antibiotics were added at following concentrations: Ampicillin 50 µg/ml, chloramphenicol 10 µg/ml, erythromycin 5 µg/ml, penicillin G (PenG) 200 µg/ml.

### DNA manipulation and cloning procedures

Isolation of plasmid DNA and transformation into *E. coli* or Rev2 were conducted according to standard procedures^26,43,44^. For subcloning of plasmid pET302/*ply006* purified pET302 vector (Invitrogen) and codon optimized synthetic DNA (Gene Art DNA Strings, Thermo Fischer Scientific) encoding *ply006* gene and appropriate restriction sites were digested using restriction enzymes NdeI and BamHI-HF (New England Biolabs) followed by ligation with T4 DNA Ligase (Thermo Fischer Scientific) and transformation into *E. coli* BL21-Gold (DE 3). Sequence identity was confirmed by Sanger sequencing (Microsynth). For subcloning of pET21a/*ply007*, the backbone of pET21a (EMD Biosciences) was amplified using primers JPR1168 and JPR1169. *Ply007* was amplified using primers JPR1170 and JPR1171. Individual fragments were assembled by Gibson assembly at 50°C for 1 h in a total reaction volume of 20 µl (NEBuilder HiFi DNA Assembly Cloning Kit, New England Biolabs), thereby adding 6x His-tag coding sequences to the 3’ end of *ply007*. Resources used in this study are disclosed in Table S1 and S 2.

### Phage propagation and purification

Soft agar overlay method was employed for phage propagation using LC Soft agar (0.4% LB agar; 10 mM MgSO_4_; 10 g/l glucose; supplemented with 10 mM CaCl_2_) as top agar and 0.5 BHI agar for plating. Phages and propagation hosts are listed in Table S1. For extraction, semi-confluent plates were incubated with 3 ml SM buffer (100 mM NaCl, 8 mM MgSO_4_, and 50 mM Tris, pH 7.4) followed by 0.2 µm sterile filtration of the suspension. All crude lysates were treated with DNAse I (10 µg/ml) and RNAse (1U/ 10ml) for 1h at 37°C. For precipitation, one volume of precipitation solution (polyethylene glycol (PEG), 3M NaCl, 30% PEG8000) was added to two volumes of lysate and incubated on ice for 24 h followed by centrifugation at 10,000xg for 15 min at 4°C. Pellets were resuspended in 5 ml SM buffer and purified via CsCl density gradient ultracentrifugation (Optima XPN-80 ultracentrifuge; Beckman Coulter) at 19,200xg for 18 h at 10°C. Purified phage bands were carefully isolated using a syringe, dialyzed two times against 1000x fold excess of SM buffer and stored at 4°C.

### Assembly of synthetic genomes, L-form transformation and genome rebooting

Assembly, transformation and rebooting of synthetic bacteriophage genomes was performed as described earlier^22^ with slight modifications. Bacteriophage genomes were designed *in silico*; resources are listed in Table S3. Briefly, codon optimized e*gfp* and a strong ribosomal binding site (RBS, GAGGAGGTAAATATAT) sequence were inserted downstream of gene *cps* (*gp07)*. Designed fragments were PCR-amplified from purified phage A006 or synthetic DNA to yield a total of six DNA fragments per phage genome followed by Gibson assembly at 50°C for 1 h in a total reaction volume of 20 µl (NEBuilder HiFi DNA Assembly Cloning Kit, New England Biolabs). Assembly reactions were carried out with purified DNA fragments to yield synthetic genomes. For L-form transfection *L. monocytogenes* Rev2 was used to for rebooting^22^. To this end, 5 µl of a frozen stock were inoculated in DM3 medium and incubated statically at 32°C for 24 h. The culture was adjusted to OD(600 nm) 0.15. For L-form transfection, 100 µl of adjusted L-form culture was mixed thoroughly with 150 µl heat-sterilized PEG8000 40% and 20 µl of Gibson assembly reaction in a 50 ml falcon tubes using wide bore pipet tips. After 5 min, 10 ml of prewarmed DM3 medium was added to the mix and incubated at 32°C for 8 h. Matured phage particles were detected by soft agar overlay method followed by screening for plaques. To this end, 5 ml of molten LC soft agar were mixed with 50 µl of transfected L-forms and 200 µl of an EGD-e overnight culture and plated on 0.5 BHI agar plates and incubated at room temperature. Individual plaques were picked after 24 h and propagated three times. For visualization, plates were scanned in transillumination mode (Image Scanner, Amersham Biosciences); contrast was adjusted for clarity where necessary.

### Time course turbidity or fluorescence assays

Time course turbidity assays were performed for wild-type phage A006 and A006::*egfp*_*cps*_ to demonstrate that the lysis kinetics of both phages are comparable. To this end, mid-exponential *L. monocytogenes* Rev2 cells expressing chromosomally integrated RFP were pelleted at 12000xg for 4 min, resuspended in DM3_Φ_ and adjusted to an OD(600 nm) of 0.1 (≈10^8^ bacteria/ml). 190 µl of diluted culture were infected with 10 µl of A006 or A006::*egfp*_*cps*_ phage lysate (10^10^ PFU/ml). Turbidity was monitored in 2 min intervals at 30 °C in flat-bottom 96 well plates using a FLUOstar OMEGA plate reader (BMG LABTECH). Plates were agitated before each measurement. Identical infections conditions were used for fluorescence time course assays. Fluorescence intensities were measured in black walled 96-wells with a FLUOstar OMEGA plate reader (BMG LABTECH) at 485 nm excitation wavelength with a 520 nm emission filter. Fluorescence time course assays were background corrected by subtraction of controls (bacteria+ phage A006). All data were acquired in three independent experiments.

### Phage adsorption assay

To quantify A006 phage adsorption to the bacterial surface, overnight cultures of *L. monocytogenes* EGD-e or mutants EGD-e *Δlmo1083* (rhamnose-deficient) or EGD-e *Δlmo2550* (GlcNAc-deficient) phage pulldown assays were performed as previously described^2^. The number of adsorbed phage particles was determined by plaque assays using the soft agar overlay method.

### Phage survival assay

To quantify L-form induction and survival in response to phage infection, overnight cultures were diluted 1:20 with 0.5 BHI or BHI-FC and grown to mid-exponential phase. Bacteria were pelleted at 12000xg for 4 min and resuspended in DM3_Φ_ medium or sterile-filtered human urine and adjusted to an OD(600 nm) of 0.0375. Decimal serial dilutions of purified phage were prepared, and 10 µl of each dilution were added to 190 µl of cell suspension followed by incubation in flat-bottom 96 wells at 30°C using a FLUOstar OMEGA plate reader (BMG LABTECH). To follow phage induced bacterial lysis over time, OD(600 nm) was monitored in 5 min intervals, plates were agitated prior to each measurement. To quantify L-form survival, serial dilutions of individual infections were plated on osmoprotective agar. Viable L-form and walled bacterial counts were enumerated 2-5 d post infection. Data were acquired in three independent experiments and three technical replicates per experiment. Ability of L-form colonies to revert in the absence of phage was tested by picking and inoculation of L-form cells on DM3 agar. Reversion (i.e., occurrence of walled cells) was confirmed by light-microscopy.

### Endolysin overexpression and purification

Codon optimized endolysin Ply006 was expressed from vector pET302; C-terminally 6xHis-tagged Ply007 was expressed from vector pet21a(+) in *E. coli* BL21 Gold (DE3) cells in LB-PE medium (15 g/L Tryptone, 8 g/L Yeast extract, 5 g/L NaCl, pH 7.8)^45^. Recombinant protein expression was induced with 0.5 mM isopropyl-β-D-1-thiogalactopyranoside (IPTG) at mid-exponential phase and allowed to proceed for 18 h at 19°C. Bacteria were harvested by centrifugation at 7,000xg for 10 min at 4°C and lysed in buffer A (20 mM Na_2_HPO_4_, 30% glycerol, pH 7.4) using a Stansted Fluid Power pressure cell homogenizer (100 MPa) followed by centrifugation at 20,000xg for 60 min at 4°C to remove cellular debris. Cleared lysates containing proteins with no His tag were purified by cation exchange chromatography (CIEX) using a 5 ml HiTrap Sepharose SP FF column (GE Healthcare) fitted on an ÄKTA fast protein liquid chromatography (FPLC) device (GE Healthcare). After washing, bound proteins were eluted with buffer B (20 mM Na_2_HPO_4_,1M NaCl, 10% glycerol, pH 7.4). 6xHis tagged protein were purified by immobilized metal ion chromatography using nickel-NTA super flow resin (Qiagen) as previously described with slight modifications^46^. Briefly, Column was washed with 25 column volumes (CV) of lysis buffer (50 mM Na_2_HPO_4_, 300 mM NaCl, 10 mM imidazole, 30 % glycerol, pH 8.0), followed by elution of target proteins with elution buffer (50 mM Na_2_HPO_4_, 300 mM NaCl, 250 mM Imidazole, 30 % Glycerol, pH 8.0) in 1 ml fractions. All purified proteins were dialyzed against 1000x excess of dialysis buffer (30% glycerol, 50 mM NaH_2_PO_4_, 300 mM NaCl, pH 7.5). Protein identity was confirmed by SDS PAGE using Mini-Protean TGX-stain free precast gels (Bio-Rad) concentration was measured using a Nanodrop ND-1000 Spectrophotometer (Thermo Fischer Scientific).

### Endolysin catalytic activity and L-form survival assay

To assess the specific activity of Ply006 on *L. monocytogenes* strain Rev2 and Ply007 on *E. faecalis* turbidity reduction of bacterial substrate cells was measured at 600 nm in flat-bottom 96-well plates using a FLUOstar OMEGA plate reader (BMG LABTECH). Briefly, cultures of *L. monocytogenes* strain Rev2 or *E. faecalis* were diluted to OD(600 nm) of 0.1 in 0.5 BHI or BHI-FC, respectively and incubated until reaching mid-exponential phase. Cells were pelleted by centrifugation at 8000xg for 5 min and resuspended in DM3_Φ_ or DM3_Φ_ supplemented with 3.2 mM L-cysteine, respectively to reach a final OD(600 nm) of 2. To determine the linear activity range, 2x serial dilutions of purified endolysin were prepared and 100 µl of each dilution were mixed with 100 µl of the corresponding cell suspension. Turbidity reduction was monitored in 5 min intervals at 30 °C for 40 min. Plates were agitated prior to each measurement, lysis curves were blank corrected against medium without endolysin and bacteria. To determine specific enzyme activities, lysis curves were fitted to a 5-parametric sigmoidal function using SigmaPlot 13 (Systat Software) as described previously^47^. The steepest slopes of individual lysis curves within the linear activity range were used to calculate the specific activities in excel (Microsoft) as described earlier^46^. All Data were acquired in three independent experiments from technical triplicates.

### Mass Spectrometry

Protein masses were identified using Liquid Chromatography-Electrospray Ionization-Mass Spectrometry (LC-ESI-MS) at the Functional Genomics Center Zürich, Switzerland (www.FGCZ.ch), using standard protocols. Briefly, prior to ESI-MS analysis, the sample was desalted using a C4 ZipTip (Millipore) and analyzed in MeOH:2-PrOH:0.2% FA (30:20:50). The solution was infused through a fused silica capillary, ID 75 μm at a flow rate of 1 μL/min and sprayed through a PicoTip, ID 30 μm (New Objective). Nano ESI-MS analysis of the samples was performed on a Synapt G2_Si mass spectrometer and the data were recorded with the MassLynx 4.2 Software (Waters). Mass spectra were acquired in the positive-ion mode by scanning an m/z range from 400 to 4000 da with a scan duration of 1 sec and an interscan delay of 0.1 sec. The spray voltage was set to 3 kV, the cone voltage to 50V, and source temperature 80°C. For every detected species, the recorded m/z data were individually deconvoluted into mass spectra by applying the maximum entropy algorithm MaxEnt1 (MaxLynx) with a resolution of the output mass 0.5 Da/channel and Uniform Gaussian Damage Model at the half height of 0.7 Da.

### Microscopic Imaging

Light microscopy and confocal laser scanning microscopy (CLSM) was performed using an inverted Leica TCS SPE research microscope (Leica Microsystems GmbH) with a HCX PL FLUOTAR 100.0 × 1.30oil objective, DFC360 FX camera and Leica application suite software v. 2.5.1.6757 fitted with an environmental chamber. For snapshot live-cell imaging, *L. monocytogenes* samples were mounted on microscopic slides covered with 1% 0.5 BHI agar or 1% DM3 agar for L-forms; *E. faecalis* samples were mounted on 1% BHI-FC agar or 1% DM3 agar supplemented with 3.2 mM L-cysteine for L-forms. To avoid drying of the agar film cover slips were sealed using transparent nail polish. All time-lapse imaging was performed at 30°C. Where appropriate, fluorescence channels were included using an excitation wavelength of 488 nm for eGFP expressing samples and 532 nm for RFP expressing samples. For time-lapse imaging of A006 ΔLCR-mediated L-form switching, exponential cultures of Rev2 cells expressing chromosomally integrated eGFP were pelleted and OD was adjusted to 0.0375 (≈3.75*10^7^ bacteria/ml) with DM3_Φ_. 190 µl of diluted culture was infected with 10 µl of A006 ΔLCR phage lysate (5×10^5^ PFU/ml) at 30°C. Time-lapse imaging was started 6 h post infection. To observe L-form proliferation time-lapse imaging was started 18 h post infection. For snapshot imaging of A006-mediated effects on L-form switching Rev2 cells were pelleted, and OD(600 nm) was adjusted to 0.0375 using DM3_Φ_ or 0.5 BHI respectively, followed by infection with 10 µl of A006 phage lysate (5×10^5^ PFU/ml or 5×10^6^ PFU/ml) at 30°C. Bacterial growth and lysis was monitored spectrophotometrically as described above and samples were imaged at several timepoints throughout the infection process. For snapshot imaging of L-forms in presence of phage, Rev2 L-form cultures expressing RFP were adjusted to an OD(600 nm) of 0.2. 190 µl of bacterial culture were mixed with 10 µl of A006::egfp_cps_ (10^10^ PFU/ml). Samples were imaged after 0, 45, 60, 75 or 120 min. For time-lapse-imaging of endolysin-treated bacteria under hypotonic or osmoprotective conditions, mid-exponential bacterial culture was pelleted and resuspended with appropriate purified endolysin to reach a final concentration of 1024 nM and an OD(600 nm) of 1. Samples were immediately mounted for microscopy. To observe endolysin-induced L-form emergence, bacteria were exposed to endolysin for 1 h at 30°C followed by time-lapse imaging. Low-magnification imaging of bacterial colonies was performed using a Leica S6 D stereo microscope equipped with a MC 170 HD camera. Image analysis and processing was performed using Fiji v.1.51 (National Institutes of Health).

### Plunge freezing

Plunge freezing was performed using a FEI Vitrobot (Thermo Fisher Scientific)^48^. For cryoET sample preparation of bacterial cells, 10 nm colloidal gold fiducial markers (Sigma-Aldrich) were added to each sample in a ratio of 1:5 (v/v) to allow tilt image alignments. For Vitrobot setup, a filter paper (Whatman, 47 mm diameter) and a Teflon sheet was installed for single sided blotting in a pre-cooled chamber (4° C) with 100% humidity. EM grids (R2/2, Cu 200 mesh; Quantifoil Micro Tools GmbH) were glow-discharged for 45 sec at 25 mA by PELCO easiGlow discharger. 4 μl sample aliquots were applied to each grid, incubated for 15 sec and blotted for 6.5 sec followed by immediate plunge freezing in an ethane-propane mixture (37% v/v ethane/ 63% v/v propane)^49^. Grids were stored in liquid nitrogen. Prior to loading of the samples into the cryo-electron microscope, the grids were clipped. For sample preparation all bacterial samples were pelleted, and OD(600 nm) was adjusted to 2-2.5 prior to blotting. If required, *L. monocytogenes* or *E. faecalis* cells were exposed to 1024 nM purified Ply006 or Ply007, respectively, followed by plunge freezing at the desired timepoints. For imaging of phage adsorption, bacterial cultures were adjusted to an OD(600 nm) of 0.1. 95 µl of diluted culture were mixed with 5 µl of purified phage lysate (10^11^ PFU/ml) bacterial cells were infected with followed by 5min incubation at room temperature.

### Cryo-electron tomography (cryoET)

For cryoET imaging, all tilt series images were collected in a Titan Krios 300 kV transmission electron microscope (Thermo Fisher Scientific) equipped with a field emission gun (FEG), an energy filter (slit width 20 eV; Gatan Inc.) and K2 or K3 direct electron detectors (Gatan Inc). Images were recorded at a pixel size of 4.34, 2.75 or 3.4 Å. Tilt series were collected from -60° to +60°degree with 2° increments and a defocus of -9 μm. A cumulative total dose of 120-150 e^-^/Å^2^ was used for acquisition. Tilt series and 2D projections images were acquired using SerialEM.

### Tomogram reconstruction

Tilt series and 2D images were automatically acquired using SerialEM^50^. Drift-correction and exposure-filtering was conducted using Alignframes. Three-dimensional reconstructions and segmentations were calculated using IMOD software package^51,52^; where appropriate deconvolution filtering was employed. Visualization and two-dimensional slices through a three-dimensional volume were acquired using 3dmod.

### Flow cytometry analysis

Flow cytometry was performed on a BD FACS Aria III cell sorting device equipped with BD FACS Diva 8.01 software (BD Biosciences). Prior to experiments, voltage settings for the relevant fluorescence channels were adjusted by running *L. monocytogenes* strain Rev2 walled cells expressing no fluorescent proteins or eGFP or RFP. Fluorescence was measured after excitation at 488 nm (eGFP) or 561nm (RFP) using 530/30 nm and 610/20 nm bandpass filter, respectively. Forward scatter (FSC-H) and side scatter (SSC-H) threshold values were set to 500 to minimize noise. For all experiments, bacterial cells expressing chromosomally integrated RFP were used. Bacterial events were identified based on scatter-(FSC-h) and RFP fluorescence intensity. To eliminate doublets, serial dilutions of bacteria were run to determine the linear range of the event rate. Flow cytometry -grade PBS, pH7.4 (Thermo Fischer Scientific) was used as sheath fluid. For analysis of phage-induced eGFP fluorescence, mid-exponential *L. monocytogenes* strain Rev2 cells expressing RFP cells were diluted to an OD(600 nm) of 0.1. 190 µl of the diluted sample were infected with 10 µl A006::*egfp*_*cps*_ 10^8^ PFU at 30°C. Samples were incubated for 45, 60, 75 or 120 min and diluted 1:50 in flow cytometry grade PBS, pH7.4. Diluted samples were immediately analysed from a 1.5 ml tube with no swirling at 4°C. The flow was adjusted to the lowest flow rate (ca. 12 μl/min) resulting in 200-500 events/sec. For each sample 10,000 events were measured. All FACS analysis was complemented by simultaneous microscopic analysis of each sample (see section Microscopic Imaging). Selected samples were chosen for reanalysis as a quality control. To this end, cell sorting was performed using a 70-micron nozzle at 87 kHz. The drop delay was set manually using BD FACS™ Accudrop Beads (BD Biosciences) before the experiment. Samples were acquired at the lowest flow rate, resulting in approximately 200-500 event/sec and reanalyzed with a target value of >95% of positive cells. Samples were collected in a tube containing 50 µl DM3_Φ_ to avoid cell damage during the collection process. All data analysis was done using FlowJo v10.6.1 (BD Biosciences).

### Data Analysis and visualization

Data analysis and plotting of data was performed in Prism v.8.0 (Graphpad Software), with the exception of FACS data, which were analyzed in FlowJo v10.6.1 (BD Biosciences).

## Data availability

All data that support the findings of this study are available from the authors on reasonable request.

## Acknowledgements

The authors would like to thank M. Wickert from the Cytometry Facility at UZH Zürich for technical support with FACS, and S. Chesnov from the Functional Genomics Center Zürich for mass spectrometry analysis of Ply007. The imaging platform ScopeM is acknowledged for instrument access. We are grateful to S. Meile for providing the *E. faecalis* Rev strain, and P. Studer for advice during preparation of the grant proposal. This work was supported by the Swiss National Science Foundation (SNSF) Grant 31003A_170042 (to M.J.L).

## Contributions

M.S. and M.J.L planned the project. J.C.W, M.J.L., M.F. and S.K. designed the research. J.C.W, M.F., A.S. and M.B. performed the experiments. J.W, A.S., M.F. and M.B analysed the data. J.W. wrote the manuscript with input from all authors. M.J.L., M.S and M.P supervised the project.

