## Supplemental Figs and Tables for "Gram-positive bacteria evade phage predation through endolysin-mediated L-form conversion"

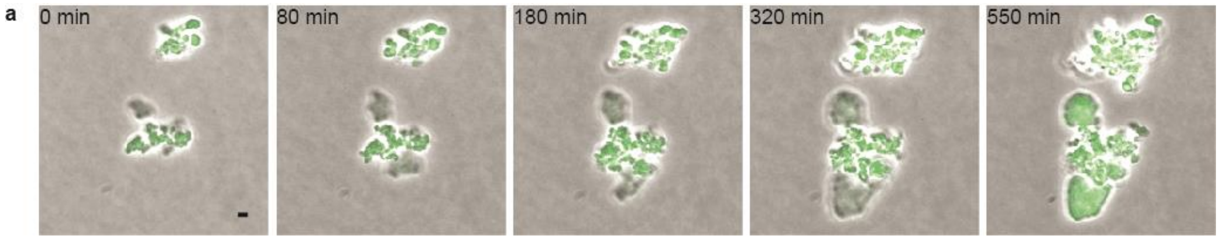


**Supplementary Figure 1: Phage infection can trigger L-form switching and proliferation.** a, Time-lapse microscopy of proliferating *L. monocytogenes* Rev2 L-forms after L-form switching in response to infection with strictly lytic phage A006 ΔLCR in $DM3_{\Phi}$ medium. Micrograph composites were obtained by merging the PC channel and the channel for green light emission. Individual Frames are extracted from Supplementary Video 2. Scale bar, 2 µm.

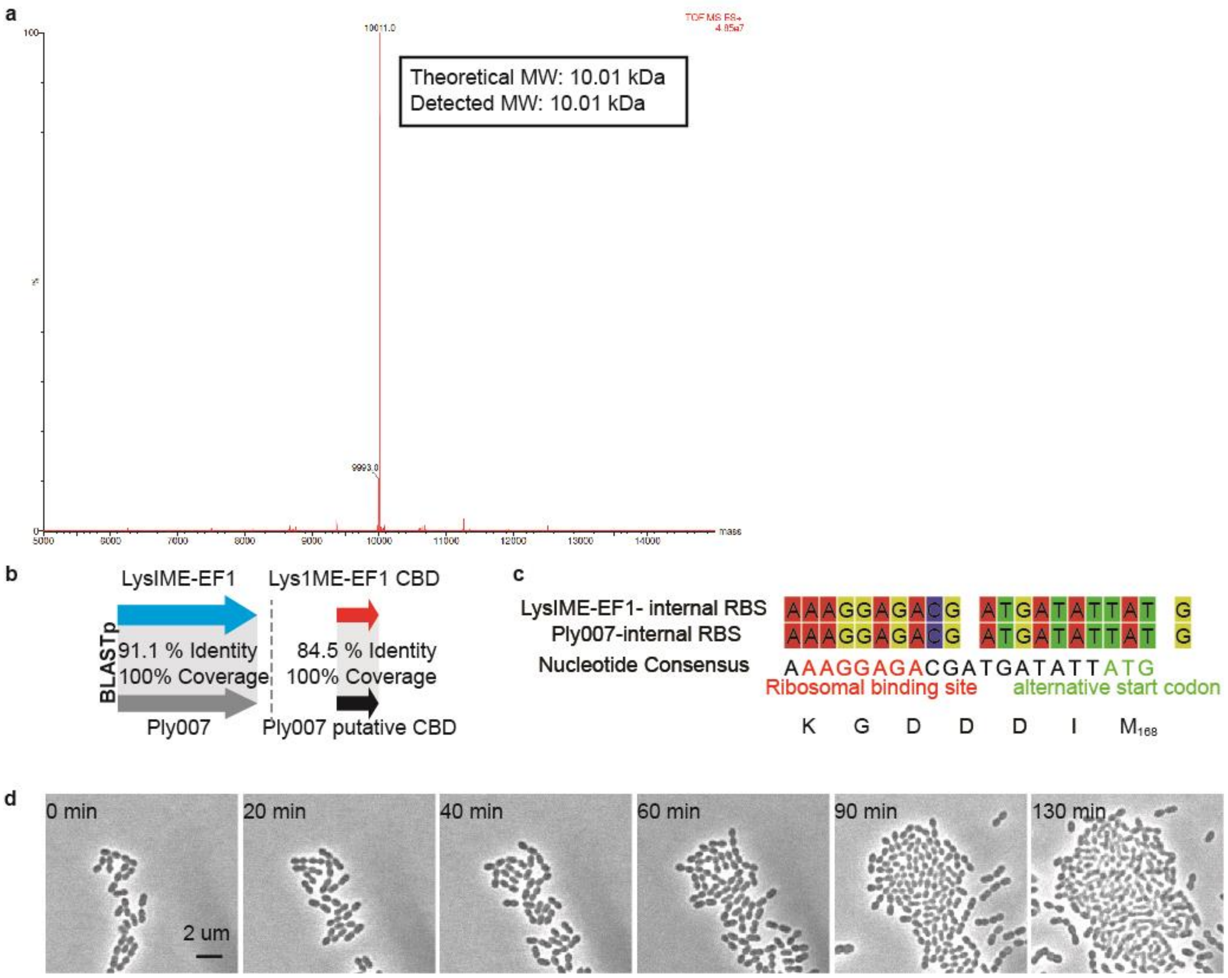


**Supplementary Figure 2: L-form switching is promoted by phage encoded endolysins under osmoprotective conditions.** a, A secondary translational product is expressed from ply007. ESI-LC-MS spectra of copurified putative Ply007 peptide fragment (secondary translational product) with fused C-terminal 6xHis-Tag; MW (fragment+Linker+6His-Tag): 10.01 kDa; peptide only:8.2 kDa. b-c, Ply007 CBD putatively forms a multimeric endolysin together with the corresponding lage subunit. b, BLASTp[53] pairwise comparison of Ply007 vs well-characterized LysIME-EF1[54] and Ply007 putative CBD vs LysIME-EF1 CBD. Shown are identity and coverage of gene products (BLASTp) in %. c, Pairwise alignment of the Lys1ME-EF1 internal ribosomal binding site region (RBS) and corresponding region in Ply007. d, Proliferation of walled E. faecalis cells on 0.5 BHI-FC agar. See also Supplementary Video 7. Scale bars; 2 µm.

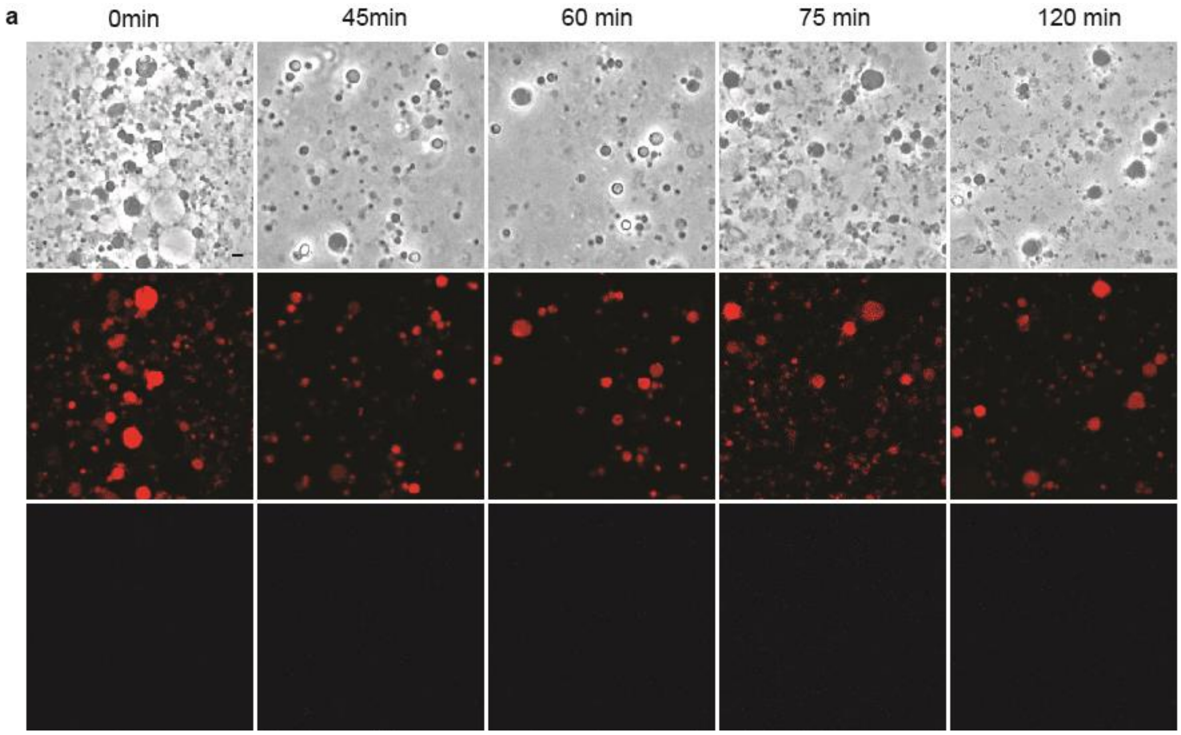


**Supplementary Figure 3: Effect of L-form exposure to engineered reporter phages** . Rev2 L-forms expressing chromosomally integrated RFP were challenged with excess amounts of phage A006::*egfp*$_{cps}$ (see also Fig. 5g). Micrographs show samples analyzed at different timepoints from 0-120 min. eGFP expression was used as a marker to screen for potential phage infection. Note, that no eGFP signal was detectable. PC micrographs and corresponding channels for red (middle panel) and green (bottom panel) light emission are shown. Scale bar, 2 µm.

**Supplementary Table 1**: Strains, phages and plasmids used in this study

| **Strains, phages, plasmids** | **purpose/remarks** | **source/reference** |
|---|---|---|
| *L. monocytogenes EGD-e* | - | Lab strain collection, SV 1/2 |
| *L. monocytogenes EGD-e Δlmo1083* | mutant lacking rhamnosylated WTAs | lab strain collection, SV 1/2a |
| *L. monocytogenes EGD-e Δlmo2550* | mutant lacking GlcNAcylated WTAs | lab strain collection, SV 1/2a |
| *L. monocytogenes Rev2* | - | *Kilcher et al.*, SV 1/2a |
| *L. monocytogenes Rev2::pPL3/egfp* | eGFP expression | this work |
| *L. monocytogenes Rev2::pPL2/rfp* | tagRFP expression | this work |
| *L. monocytogenes Rev2-P100 ("1005")* | phage rebooting | this work |
| *L. monocytogenes Mack* | propagation host for ΦP35 | Lab strain collection, SV 1/2 |
| *L. monocytogenes WSLC 1001* | propagation host for ΦA118 | lab strain collection, SV 1/2 |
| *L. ivanovii* WSLC3009 | propagation host for ΦA511, ΦP40 | lab strain collection, SV 5 |
| *E. coli* BL21 Gold (DE3) | protein expression | Stratagene |
| *E. coli* XL1-Blue MRF' | cloning | Stratagene |
| *E. coli* XL1-blue MRF' pPL3/*egfp* | plasmid amplification | *Studer et al.*, 2016 |
| *E. coli* XL1-blue MRF' pPL2/*rfp* | plasmid amplification | *Studer et al.*, 2016 |
| *E. faecalis* ATCC 19433 | - | lab strain collection |
| *E. faecalis* Rev | - | this work |
| ΦA006 | - | lab phage collection |
| ΦA006 ΔLCR | Lysogeny control region deleted --> strictly lytic | *Meile et al.*, 2020 |
| ΦA006::$eGFP_{cps}$ | eGFP expression | this work |
| ΦA118 | - | lab phage collection |
| ΦA511 | - | lab phage collection |
| ΦP35 | - | lab phage collection |
| ΦP40 | - | lab phage collection |
| ΦEfs7 | - | lab phage collection |
| pPL2/*rfp* | Cam$^R$ (gram-, gram+), RFP expression | *Studer et al.*, 2016 |
| pPL3/*egfp* | Cam$^R$ (gram-), EryR (gram+), eGFP expression | *Studer et al.*, 2016 |
| pET302 | Amp$^R$, for protein expression | this work |
| pET302/*ply006* | Amp$^R$, PlyA006 expression, based on pET 302 | this work |
| pET21a | Amp$^R$, for protein expression | lab stock |
| pET21a/*ply007* | Amp$^R$, PlyA007 expression, based on pET21a | this work |

*Antibiotic resistances: Cam$^R$: chloramphenicol, Amp$^R$: ampicillin, Ery$^R$: erythromycin

**Supplementary Table 1**: Sequencing oligonucleotides and DNA strings used in this study.

| ID | Sequence (5'-3') | purpose |
|---|---|---|
| T7-1 | AATACGACTCACTATAGGG | sequencing primer pET302 |
| pET-RP | CTAGTTATTGCTCAGCGG | sequencing primer pET302, pET21a |
| T7 | GAAATTAATACGACTCACTATAGGG | sequencing primer pET21a |
| JPR1168 | TTTAAGAAGGAGATATACATATGGTTAAAGTAAACGATGTAG | pET21a backbone amplification, assembly pET21a/ply007 |
| JPR1169 | GCGGCCGCAAGCTTGTCGACTAACTTAACTTGTGGGTAAGC | pET21a backbone amplification, assembly pET21a/ply007 |
| JPR1170 | GTCGACAAGCTTGCG | *ply007* amplification, assembly pET21a/ply007 |
| JPR1171 | ATGTATATCTCCTTCTTAAAGTTAAAC | *ply007* amplification, assembly pET21a/ply007 |
| String A (*eGFP*, codonoptimized*)* | ATGTCTAAAGGTGAAGAATTATTCACTGGTGTTGTTCCAATCTTAGTTGAATTAGATGGTGATGTTAACGGTCATAAATTCTCTGTTTCTGGTGAAGGTGAAGGTGATGCTACTTACGGTAAATTAACTTTAAAATTCATCTGTACTACTGGTAAATTACCAGTTCCATGGCCAACTTTAGTTACTACTTTCGCTTACGGTTTACAATGTTTCGCTCGTTACCCAGATCATATGAAACAACATGATTTCTTCAAATCTGCTATGCCAGAAGGTTACGTTCAAGAACGTACTATCTTCTTCAAAGATGATGGTAACTACAAAACTCGTGCTGAAGTTAAATTCGAAGGTGATACTTTAGTTAACCGTATCGAATTAAAAGGTATCGATTTCAAAGAAGATGGTAACATCTTAGGTCATAAATTAGAATACAACTACAACTCTCATAACGTTTACATCATGGCTGATAAACAAAAAAACGGTATCAAAGTTAACTTCAAAATTAGACACAACATTGAAGATGGAAGCGTTCAACTAGCAGACCATTATCAACAAAATACTCCAATTGGCGATGGCCCAGTTTTATTACCAGATAACCATTACTTATCTACTCAATCTGCTTTATCTAAAGATCCAAACGAAAAACGTGATCATATGGTTTTATTAGAATTCGTTACTGCTGCTGGTATCACTCATGGTATGGATGAATTATACAAATAA | construction of phage $\Phi$A006::*eGFP*$_{cps}$ |

| | | |
|---|---|---|
| String B (*ply006*, codonoptimized + *BamHI/NdeI* restriction site) | TATACATATGGCACTGACCGAAGCATGGCTGATTGA<br>AAAAGCAAATCGTAAACTGAATGTGAGCGGCATGAA<br>TAAAAGCGTTGCAGATAAAACCCGCAACGTGATCAA<br>AAAAATGGCCAAAAAAGGCATCTATCTGTGTGTTGCA<br>CAGGGTTATCGTAGCAGCGCAGAACAGAATGCACTG<br>TATGCCCAGGGTCGTACCAAACCGGGTGCAGTTGTT<br>ACCAATGCAAAAGGTGGTCAGAGCAATCATAACTAT<br>GGTGTTGCAGTTGATCTGTGCCTGTATACCAGTGAT<br>GGTAAAAATGTTATTTGGGAAAGCACCACCAGTCGTT<br>GGAAAACCGTTGTTAGCGCAATGAAAGCCGAAGGTT<br>TTGAATGGGGTGGTGATTGGAAAAGCTTTAAAGATTA<br>TCCGCACTTCGAACTGTATGATGCAGCCGGTGGCGA<br>AAAAGCACCGAGCACCAGCGCAAGCAAACCGGCAA<br>CCAGCACCAGCAGTAACAAAAATGTGTATTACACCG<br>AGAATCCGCGTAAAGTTAAAACCCTGGTTCAGTGCG<br>ATCTGTATAATAGCGTTGATTTTACCGAGAAACATAA<br>AACCGGTGGCACCTATCCGGCAGGCACCGTGTTTAC<br>CATTAGCGGTATGGGTAAAACCAAAGGTGGTACACC<br>GCGTCTGAAAACCAAAAGCGGTTATTATCTGACCGC<br>CAACAAAAAGTTCGTGAAAAAAATCTAAGGATCCGGC<br>T | construction of pET302/*ply006* |

*Underlined: coding DNA sequences

**Supplemenentary Table 3**: Synthetic phage ΦA006::*egfp*$_{cps}$ assembly: Oligonucleotides, templates, and DNA fragments

| Fragment ID | Template | ID primer fwd | Sequence (5'-3') | ID primer rev | Sequence (5'-3') | fragment size (nt) |
|---|---|---|---|---|---|---|
| f1 | synthetic DNA | JPR668[S] | TCTACAGAAGTCTAAG**AGGAGG**TAAATATATATGTCTAAAGG | JPR669[S] | ATTTCCCTAACCTCCTTATTTGTATAATTCATCCATACCATG | 763 |
| f2 | A006 gDNA | JPR670 | AAGGAGGTTAGGGAAATGCAATTAAAAAAAGAAAATGTCGTTTAC | SK684 | GAATATCCTAGCGAATGCGAAATAG | 3826 |
| f3 | A006 gDNA | SK683 | TGAACTATGTCGGTCCTATTTCG | SK686 | AATAAATACAATACTTACACCTGGAATG | 9294 |
| f4 | A006 gDNA | SK685 | ATTACAATAGGCCATTCCAGGTG | SK688 | CCAAAGATCATCAACAACCATCG | 9711 |
| f5 | A006 gDNA | SK687 | AAACATGGTTGAGAATCCGATGG | SK682 | CTTACTGGCAATTTTCACAAGTGG | 9294 |
| f6 | A006 gDNA | SK681 | TTCAGATAAGACAATGCCACTTGTG | JPR671 | TATATTTACCTCCTCTTAGACTTCTGTAGAAGCGATTACG | 6176 |

*Bold: RBS inserted via primer, underlined: primer binding site
[S]used for Sanger sequencing of eGFP insert
